# ESGq: Alternative Splicing events quantification across conditions based on Event Splicing Graphs

**DOI:** 10.1101/2023.07.05.547757

**Authors:** Davide Cozzi, Paola Bonizzoni, Luca Denti

## Abstract

Alternative Splicing (AS) is a regulation mechanism that contributes to protein diversity and is also associated to many diseases and tumors. Alternative splicing events quantification from RNA-Seq reads is a crucial step in understanding this complex biological mechanism. However, tools for AS events detection and quantification show inconsistent results. This reduces their reliability in fully capturing and explaining alternative splicing. We introduce ESGq, a novel approach for the quantification of AS events across conditions based on read alignment against Event Splicing Graphs. By comparing ESGq to two state-of-the-art tools on real RNA-Seq data, we validate its performance and evaluate the statistical correlation of the results. ESGq is freely available at https://github.com/AlgoLab/ESGq.

## 1. Introduction

Alternative Splicing (AS) is a post-transcription regulation mechanism that contributes to isoform and protein diversity in eukaryotes. Due to AS, depending on its environment, a single gene can produce multiple isoforms, hence complicating our understanding of the gene expression process. For instance, more than 95% of multi-exon human genes [1, 2] and more than 60% of multi-exon Drosophila Melanogaster genes [3] exhibit more than one isoform. Due to its association to aging [4], cancer [5], and neuro-degenerative diseases [6], the analysis of AS is of the utmost importance.

In the last decade, RNA-Sequencing has become the de-facto standard for the analysis of alternative splicing and a plethora of tools have been proposed in the literature. From a very high level point of view, the approaches for the analysis of alternative splicing available in the literature can be divided in two groups, depending at which level they work: transcript-based [7, 8] and event-based approaches.

In this work, we will focus on the second category. This kind of approaches characterizes AS at the most fine-grained level by giving a detailed and strict description of what happens at the exon-exon (or splice) junction level. Although being more rigorous in the description of AS events, this kind of approaches resulted more ac-curate than the transcript-based competitors [9], thus potentially providing a more detailed characterization of alternative splicing. Classical AS events are grouped in 5 categories [10]: exon skipping, alternative 3’ (acceptor) splice sites, alternative 5’ (donor) splice sites, intron retention, and mutually exclusive exons. Many tools have been developed to perform AS events detection and quantification [11, 12, 13, 14, 15]. Recent works [16, 17] argue that the classic definition of AS events is not satisfactory and not adequate to fully capture the complexity of alternative splicing. To this aim, they introduce Local Splicing Variations, a novel concept that aims to represent complex AS patterns and then increase the expressive power of the classical and more strict classification. However, the detection and quantification of classical AS events is an already hard - and not fully solved - problem that does not need further complications. Indeed, tools and methodologies show several limitations [9]. For instance, although AS events exhibit a strict definition, tools available in literature inconsistent results [9], due to the different definitions and filtering criteria adopted. Moreover, every tool uses its own format to describe the AS events, making any downstream analysis quite complex.

In this context, we focus on the detection and quantification of non-novel AS events across conditions and provide an extensive comparison of two state-of-the-art tools, rMATS [11] and SUPPA2 [12]. Moreover, to validate our findings, we introduce ESGq, a novel graph-based methodology for the quantification of AS events across two conditions. Inspired by the recent progress and development in the field of pangenomic and graph algorithms [18, 19], ESGq models AS events as local splicing graphs, called *event splicing graphs*, and then quantify the events by aligning reads to them. The usage of graphs in the transcriptomic world is not new [20, 13, 14] but the use of simplified graph-based representation characterizing precise loci of the genome is something that was never investigated.

Experiments on a very recent real RNA-Seq dataset sequenced from Drosophila Melanogaster flies at two time points show that, by pairing simple graphs with accurate mapping, ESGq is able to achieve comparable results to state-of-the-art without sacrificing its efficiency. Moreover, our comparison proves once again the inconsistency of the results obtained by different methodologies for the detection and quantification of AS events.

## 2. Method

We introduce ESGq, a novel graph-based method for the differential quantification of alternative splicing events across conditions. ESGq takes as input a reference genome (in FASTA format), a gene annotation (in GTF format), and a two conditions RNA-Seq dataset with optional replicates (in FASTQ format), and computes the differential expression of annotated AS events (in custom text format). For each event ESGq provides the Percent-Spliced In (PSI, *ψ*) with respect to each input replicate and the Δ*ψ*, summarizing the differential expression of each event across the two conditions. Current implementation focuses on four types of alternative splicing events (exon skipping, intron retention, alternative acceptor site, and alternative donor site) and supports both paired-end and single-end RNA-Seq datasets.

Differently from state-of-the-art approaches, which rely on spliced read alignment to a reference genome or quasi-mapping transcript quantification, ESGq relies on read alignment against a graph-based structure representing the events that need to be quantified. Instead of using a full splicing graph or a pantranscriptome, that are common structures in the literature [13, 14, 19], ESGq limits its computation to smaller and less complex graphs, the *Event Splicing Graphs*. An event splicing graph is a splicing graph which encodes only the exons and splice junctions involved in an alternative splicing event. Differently from splicing graphs commonly used in the literature, that represent all known transcripts of a gene, and differently from pantranscriptomes, where entire gene loci and intergenic regions are represented, an Event Splicing Graph encodes only the portions of the two transcripts involved in an event. By using this simpler representation, ESGq is able to achieve great efficiency without sacrificing its accuracy.

ESGq consists of three steps (also depicted in Figure 1):

**Figure 1:**
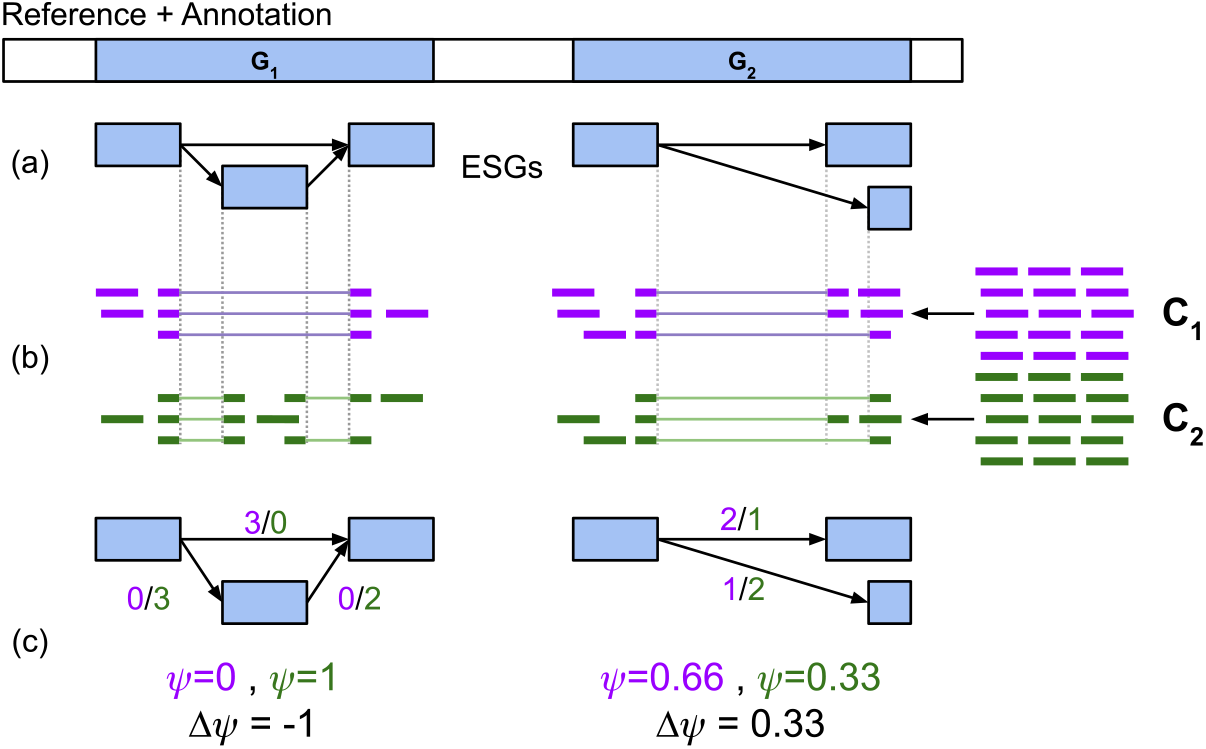
ESGq method. (a) From the reference genome and the gene annotation, ESGq builds the Event Splicing Graphs (ESGs in the figure). (b) RNA-Seq reads from the two conditions (*C*_1_ and *C*_2_) are aligned to the Event Splicing Graphs. (c) Graph alignments are used to weight junction edges and the weights are used to compute the *ψ* values (one per condition) and the Δ*ψ* value (one per dataset).

1. event splicing graphs construction
2. read alignment against event splicing graphs
3. *ψ* and Δ*ψ* computation

ESGq starts its computation by extracting the annotated alternative splicing events from the input gene annotation. To this aim, it employs a module of the SUPPA2 tool [12]. The output of this module is a list of annotated alternative splicing events, that are events whose two isoforms are already annotated in the input gene annotation. For each event, SUPPA2 reports the type (one among SE, RI, A3, A5) and the genomic coordinates of the splice junctions involved in it. Starting from this list, ESGq builds the event splicing graphs, one per event. We note that multiple graphs can be built from the same gene. By exploiting the genomic coordinates of an event, ESGq retrieves the corresponding exons and adds them as nodes in the event splicing graph. Depending on the AS event type, ESGq adds edges between these nodes in order to represent the two isoforms involved in the event: the canonical isoform (that, in graph terms, is the path denoted as 𝒫_*C*_) and the alternative one (denoted as 𝒫_*A*_). More precisely, the four scenarios, one per event type, contemplated by ESGq are depicted in Figure 2 and can be formally defined as follow:

**Figure 2:**
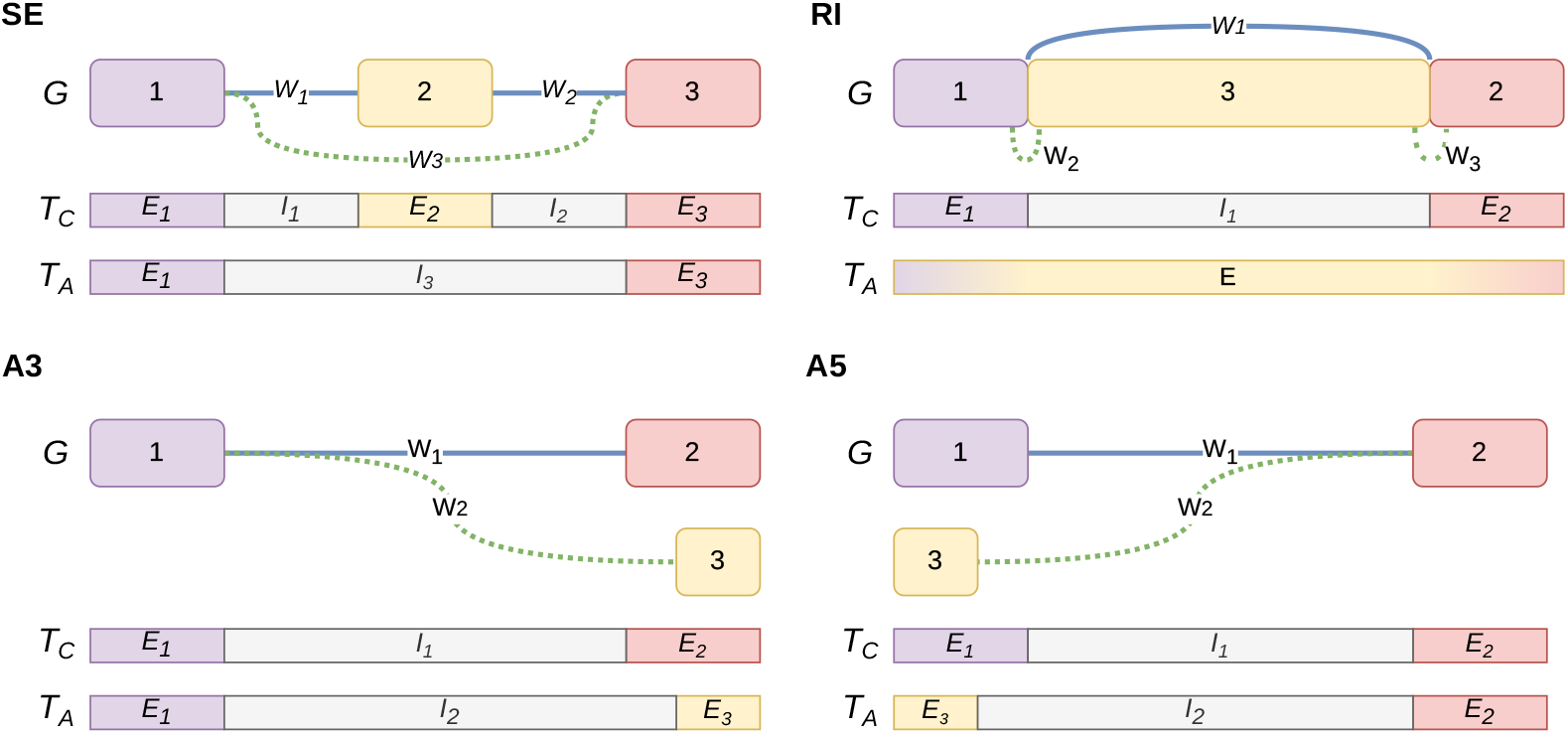
Event splicing graphs computed by ESGq. For each AS event type (exon skipping, SE, alternative acceptor site, A3, alternative donor site, A5, and intron retention, RI), we report the event splicing graph *G* and the two annotated isoforms involved in the event: the canonical transcript *T*_*C*_ (represented in *G* by blue edges) and the alternative transcript *T*_*A*_ (represented in *G* by dotted green edges). In *T*_*C*_ and *T*_*A*_, *E* blocks represent exons and *I* blocks represent introns. The edge labels *W* represent the weights computed by ESGq after the read alignment step.

- to model an exon skipping event (SE), ESGq needs to take into account three exons and this implies that the corresponding event splicing graph is composed of three nodes *n*_1_, *n*_2_, *n*_3_. In detail, *n*_2_ represents the exon that is spliced out during the event. The canonical isoform is represented by the path involving all three nodes (𝒫_*C*_ = *n*_1_ → *n*_2_ → *n*_3_) while the alternative isoform consists in the skip of *n*_2_ (𝒫_*A*_ = *n*_1_ → *n*_3_);
- to model an alternative acceptor site event (A3), ESGq needs to take into account three exons and accordingly three nodes *n*_1_, *n*_2_, *n*_3_. In detail *n*_1_ represents the upstream exon, *n*_2_ the canonical downstream exon, and *n*_3_ the downstream exon with the alternative acceptor splice site. The canonical isoform is represented by the path involving the shared upstream exon and the canonical downstream exon (𝒫_*C*_ = *n*_1_ → *n*_2_) whereas the alternative isoform changes the downstream exon with the alternative one (𝒫_*A*_ = *n*_1_ → *n*_3_);
- to model an alternative donor site event (A5), ESGq needs to take into account three exons and accordingly three nodes *n*_1_, *n*_2_, *n*_3_. In detail, *n*_1_ represents the canonical upstream exon, *n*_2_ the canonical downstream exon, and *n*_3_ the upstream exon with the alternative donor splicing site. The canonical isoform is represented by the path involving the canonical upstream exon and the shared downstream exon (𝒫_*C*_ = *n*_1_ → *n*_2_) whereas the alternative isoform changes the upstream exon with the alternative one (𝒫_*A*_ = *n*_3_ → *n*_2_);
- to model an intron retention event (RI), ESGq needs to take into account three exons. However, this case is harder than the previous ones: the three nodes *n*_1_, *n*_2_, *n*_3_ of the graph do not closely correspond to three exons, but one of them (*n*_3_) correspond to a portion of it. In detail, *n*_1_ and *n*_2_ represent the two upstream and downstream exons whereas *n*_3_ represents the retained intron (i.e., the internal portion of the exon linking the upstream and downstream canonical exons). The canonical isoform is then represented by the path involving the upstream exon and the downstream exon (𝒫_*C*_ = *n*_1_ → *n*_2_) whereas the alternative isoform includes the retained intron in the path (𝒫_*A*_ = *n*_1_ → *n*_3_ → *n*_2_).

We note that, from a conceptual point of view, each event contributes to an event splicing graph, but, from a more practical point of view, ESGq builds a single graph with multiple connected components, one per event.

In the second step, ESGq indexes the graph constructed in the previous step. Since the goal is to align the input RNA-Seq reads to the graph using the Giraffe aligner [21], ESGq employs the VG toolkit [22] to build the gBWT (graph Burrows–Wheeler Transform) index [23]. Each input replicate is then independently aligned to the graph using Giraffe. We note that, since by default vg breaks each node longer than 32bp into smaller nodes (of length ≤ 32), in order to keep the association between the nodes in the input graph (that is the one built by ESGq) and the nodes in the indexed version, ESGq directly breaks the nodes while building the event splicing graphs and links them accordingly to maintain the same two paths, i.e., isoforms. Due to this, in an event splicing graph we can observe two kind of edges: edges linking the smaller (≤ 32*bp*) nodes internal to an exon and edges that represent the real splice junction of interest for the AS event. Although ESGq differentiates between these two kinds of edge, conceptually only the edges representing a splice junction are used by ESGq. For this reason, we decided to omit these edges from Figure 2.

In the third and last step, ESGq computes the *ψ* value of the events w.r.t. each replicate and then summarize these values by comparing the two conditions and computing a Δ*ψ* value per event. This value represent the differential expression of each event across the two input conditions. To do so, ESGq analyses the graph alignments computed in the previous step and assign a weight to each edge that represents a splice junction. Since each read is aligned to a path of the graph, computing this weight is straightforward as increasing a counter per edge. Indeed, a read can be aligned to a single node of the graph, hence without using any edge, or to a sequence of nodes, hence using one or more edges. In such a case, ESGq checks every edge used by the alignment and, if an edge is a junction edge, it increases its weight by 1. In other words, since each junction edge represent a splice junction, its weight represents the number of reads that have been spliced aligned over it.

Finally, ESGq uses these weights to compute the *ψ* value of each AS event following its classical formulation, i.e., the proportion of reads supporting the standard isoform over the reads supporting both isoforms [24]. Differently from other approaches, which rely on both spliced and not spliced reads, ESGq *ψ* computation is based only on spliced reads counts, hence the support of an isoform is approximated using only these values and does not take into account its full coverage. We believe that this is a good approximation of the correct ψ value and this is also confirmed by our experimental evaluation. ψ calculation can be summarized as follows (we also refer the reader to Figure 2):

- 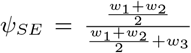 considers the mean of the weights *w*_1_, *w*_2_ of the canonical isoform and the weight *w*_3_ of the alternative isoform (in a similar fashion to [25]);
- 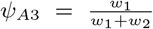 and 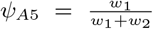 consider the weight *w*_1_ of the canonical isoform and the weight *w*_2_ of the alternative isoform
- 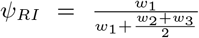 considers the weight *w*_1_ of the canonical isoform and the mean of the weights *w*_2_, *w*_3_ of the alternative isoform

Starting from these *ψ* values (one per event per replicate), which summarize the event quantification for each input replicate, ESGq computes the differential quantification across the two input conditions (Δψ) as the difference between the absolute value of the *ψ* means in the two conditions. Differently from other approaches, ESGq does not assign a *p value* to the Δψ. This is mainly a consequence of the simplified AS events quantification based only on spliced reads counts. Future works will be devoted to improve the statistical validation of ESGq results.

## 3. Experimental evaluation

We implemented ESGq in Python and the code is freely available at https://github.com/AlgoLab/ESGq. We assessed its efficacy and efficiency on a real dataset of RNA-Seq reads (SRA BioProject ID: PRJNA718442) that comes from a recent study [26] on the correlation between ageing and differential gene expression in Drosophila Melonogaster. More precisely, the study conducted a genome-wide differential expression analysis at two different time points: day 1 and day 60 flyes. The dataset consists of three replicates for two conditions (the two time points), for a total of 6 Illumina Hiseq samples. All samples are paired-end and consist of 151bp-long reads (see Table 1 for more details).

**Table 1.**
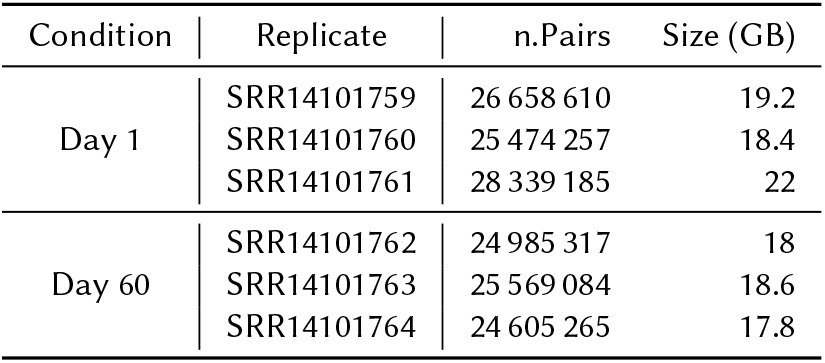
Real dataset used in our experimental evaluation (SRA Bio-Project ID: PRJNA718442). Table reports the number of reads and the size in GigaByte of each paired-end replicate.

Differently from the aforementioned study, where the focus was the analysis of differential gene expression, in this work, we analyze the same dataset from the perspective of differential quantification of alternative splicing events. To do so, we applied ESGq and two other state-of-the-art approaches for the differential quantification of alternative splicing events across multiple conditions: rMATS [11] (version 4.1.2) and SUPPA2 [12] (version 2.3). The former performs differential quantification starting from read alignment to the reference genome whereas the latter starts from the quasi-mapping transcript quantification of Salmon [27]. In this way, we have been able to compare three methodologies based on completely different frameworks: graph-based alignment, reference-based alignment, and transcript-based quasi-mapping. rMATS was run starting from the alignments produced by STAR aligner [28] (version 2.7.10b) whereas SUPPA2 was run starting from Salmon transcript quantification (version 1.10.1). All tools were run using their default parameters and 16 threads (when possible). In our analyses, we considered the reference, gene annotation, and transcripts provided by FlyBase [29], release 6.52.

We compared the Δ*ψ* values reported by the three considered tools. ESGq reported 3 276, rMATS reported 3 699, and SUPPA2 reported 1 619 AS events correctly quantified (i.e., with Δ*ψ* = *NaN*). The huge difference in the number of reported events proves the complexity of detecting AS events from RNA-Seq reads and highlights the inconsistency between different methodologies based on different filtering criteria [9]. In our analysis we included all those events correctly quantified by all three tools and considered statistically significant by both rMATS and SUPPA2, i.e., events with p value ≤ 0.05. A total of 933 events resulted from this filter: 374 exon skipping (40%), 190 alternative 3’ (20%), 154 alternative 5’ (17%), and 215 intron retention (23%). We note that we also tried to include Whippet [15] in our evaluation but, due to a different AS event representation that cannot be easily compared to the representation given by the other tools, we ended up omitting it from the analysis.

Figure 3 and Table 3 report the results of our analysis. All tools achieved comparable results. Remarkably, ESGq and rMATS achieved a very good correlation, with a Pearson correlation coefficient equal to 0.918 (Figure 3a). Although both approaches are based on read alignment, they show two main methodological differences that slightly affect their results. Firstly, rMATS uses read counts coming from read alignment to a reference genome whereas ESGq uses (spliced) read counts coming from alignment to event splicing graphs, that are a reduced representation. Secondly, the quantification step implemented in ESGq is quite simplistic and not elaborate as the probabilistic framework implemented in rMATS. However, none of these two differences (that are a wanted restriction and a current - undesired - limitation) seems to affect the results of ESGq. On the other hand, SUPPA2 resulted less correlated to the two other approaches, showing a Pearson correlation coefficient ranging from 0.766 to 0.813 (Figure 3b and 3c). This was somewhat expected since it is based on transcript quantification, and not on spliced read alignment. As proven in the literature, such a difference is expected since current transcript annotation models may result inaccurate [15].

**Figure 3:**
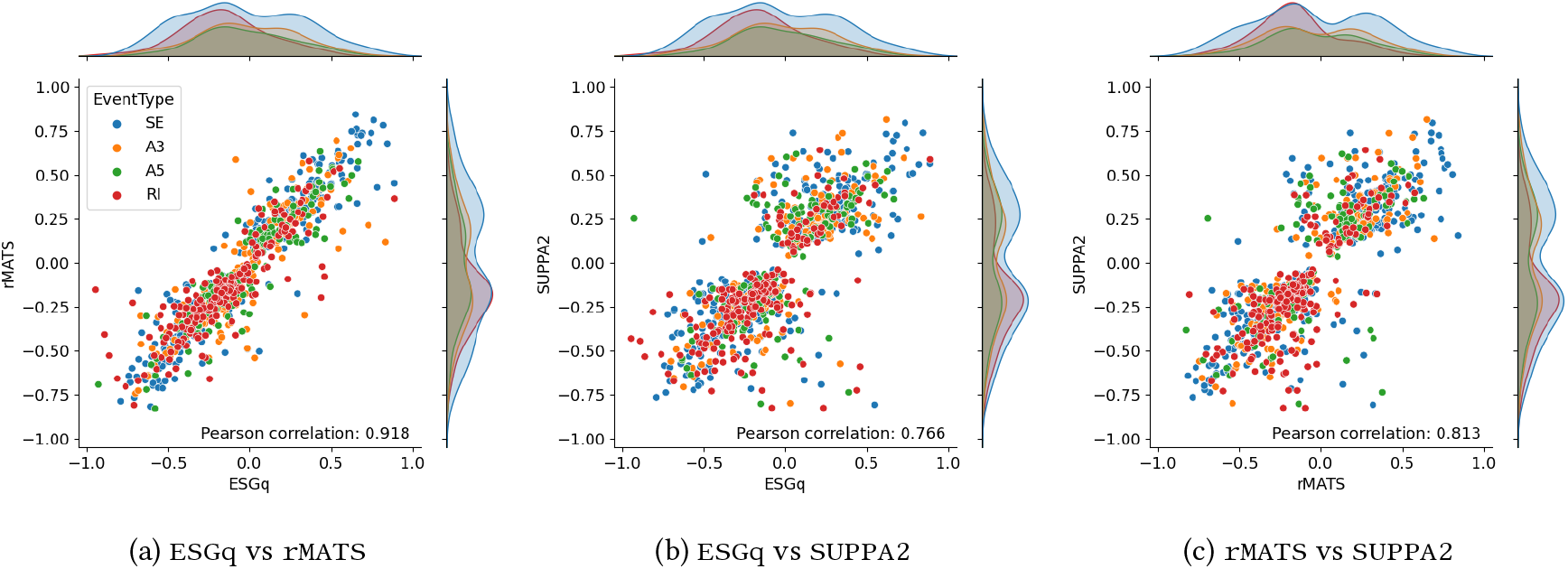
Correlation plots between the Δ*ψ* values computed by ESGq, rMATS, and SUPPA2. These results refer to our analysis of 151*bp* paired-end sample and *k* = 31 (for Salmon and SUPPA2).

Table 2 reports the correlation between the three considered tools broken down by event type. Surprisingly there is no clear trend that can be observed. ESGq and rMATS exhibit the highest correlation on exon skipping events and the lowest on intron retentions. ESGq and SUPPA2, instead, show higher correlation on exon skippings and lower correlation on alternative donor events. Finally, rMATS and SUPPA2 exhibit higher correlation on alternative acceptor site and lower correlation on alternative donor events. These results are somewhat unexpected and require a further investigation.

**Table 2.** Pearson correlation coefficients between ESGq, rMATS, and SUPPA2 (over the transcript quantification of Salmon ran with *k* = 31) broken down by event type on the 151bp paired-end dataset.

| Event type | Tool1 | Tool2 | Pearson |
| --- | --- | --- | --- |
| SE | ESGq | rMATS | 0.952 |
|  | ESGq | SUPPA2 | 0.808 |
|  | rMATS | SUPPA2 | 0.836 |
| A3 | ESGq | rMATS | 0.868 |
|  | ESGq | SUPPA2 | 0.786 |
|  | rMATS | SUPPA2 | 0.862 |
| A5 | ESGq | rMATS | 0.920 |
|  | ESGq | SUPPA2 | 0.666 |
|  | rMATS | SUPPA2 | 0.708 |
| RI | ESGq | rMATS | 0.859 |
|  | ESGq | SUPPA2 | 0.677 |
|  | rMATS | SUPPA2 | 0.760 |

**Table 3.**
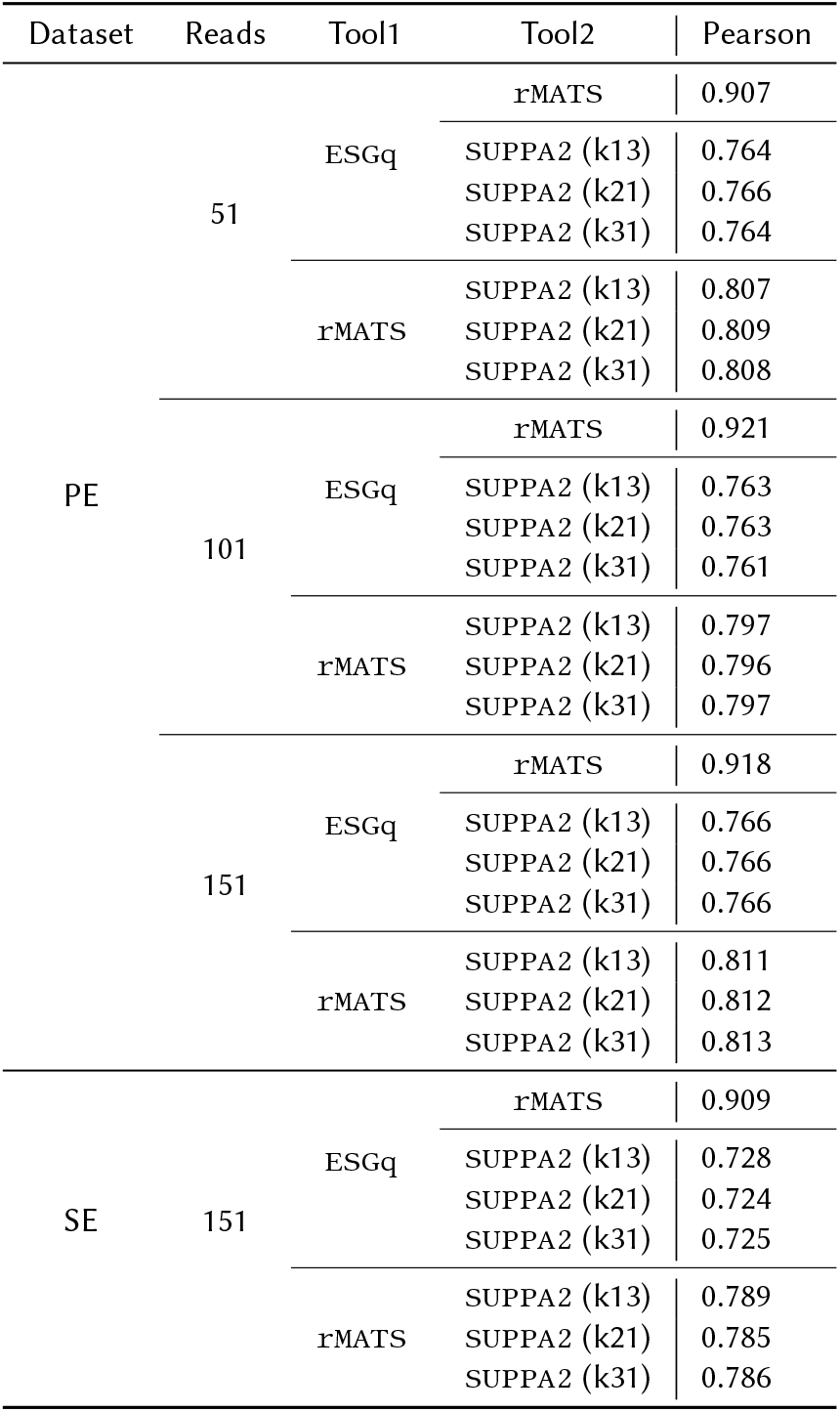
Pearson correlation coefficients between any combination of tools, i.e., ESGq, rMATS, and SUPPA2 (over the transcript quantification of Salmon ran with different *k* values) and experimental settings, i.e., different read lengths (51, 101, and 151bp) and paired/single-end dataset (PE/SE).

ESGq also resulted very computationally efficient, completing the analysis in half an hour requiring 1GB of RAM. Similarly, SUPPA2 ran in less than 10 minutes and used 1.5GB of RAM. On the contrary, rMATS resulted the most expensive approach, requiring more than 5 hours and 8GB of RAM. The most expensive step is read alignment with STAR, that required from half an hour to two hours per sample. By pairing simple and precise graph representation of well localized loci of the genome (i.e., the event splicing graphs) with fast and accurate read alignment, ESGq is able to achieve results comparable to the other alignment-based approach, while being 10x faster. Since SUPPA2 is based on the *k*-mer based quasi-mapping of Salmon, we also analyzed how *k*-mer size affects its results. For this reason, we ran Salmon (and, consequently, SUPPA2) two additional times with *k* ∈ {13, 21} (we note that 31 is the default value used in the previous results). As shown in Table 3, the results of SUPPA2 seems to be unaffected by the choice of the *k* parameter. Indeed, the correlation between SUPPA2 (ran with different *k* value) and other tools changes marginally.

Moreover, to evaluate if the results of the considered methodologies are affected by read length, starting from the 151-bp paired-end dataset, we manually trimmed the input reads to 51bp and 101bp using seqtk and created two additional datasets. Table 3 reports the results of this analysis. Even with very short reads (i.e., 51bp reads), the three tools achieved the same correlation (with a very marginal difference of < 0.015).

Finally, to evaluate how much paired-end information may improve the accuracy of the tools, we merged the two pairs of each replicate into a single sample, in order to simulate a single-end dataset. Surprisingly, there is small to none difference between the results on paired-end and single-end dataset, highlighting the robustness of the considered approaches. For instance, the Pearson correlation coefficient between ESGq and rMATS decreased by a very marginal 0.009 whereas correlation between SUPPA2 and the other approaches decreased by 0.041 (w.r.t. ESGq) and 0.027 (w.r.t. rMATS).

The experimental evaluation has been implemented as a Snakemake workflow [30], thus it is fully reproducible and easily replicable. Scripts and instructions are available at https://github.com/AlgoLab/ESGq. All the experiments were performed on a 64bit Linux (Kernel 5.15.0) system equipped with two 16-core AMD EPYC 7301 2.2GHz processors and 128GB of RAM.

## 4. Conclusions

In this paper we introduced ESGq, a novel graph-based approach for the AS event quantification across two conditions. Differently from state-of-the-art tools, ESGq is based on read alignment against local graph structures, introduced here as *event splicing graphs*, that represent AS events precisely represented in a given gene annotation. An extensive exploratory analysis on real RNA-Seq dataset showed that ESGq is able to achieve comparable results with respect to other approaches based on the alignment of reads to the reference genome, while being 10x faster.

Future works will be devoted to improving the statistical framework behind the *ψ* and Δψ computation and to extending the experimental evaluation by considering human samples, including more tools in the comparison, and evaluating the actual accuracy of ESGq. An interesting future direction consists in extending the notion of event splicing graphs to include information on known genetic variations (SNPs and indels) in order to improve the quality of read alignment, as proven in a recent work on pantrascriptomes [19]. Moreover, the detection and quantification of novel AS events from pantrancriptomes remain another interesting open problem.

## Acknowledgements

The authors thank Simone Ciccolella and Yuri Pirola for insightful discussion and technical advice.

